# Estimating Propulsion Kinetics in Absence of a Direct Measurement of the Anterior Component of Ground Reaction Force

**DOI:** 10.1101/2024.02.19.581016

**Authors:** Hannah N. Cohen, Miguel Vasquez, Fabrizio Sergi

## Abstract

Anterior ground reaction force (AGRF) is a common measurement of walking function in post-stroke individuals. It is typically measured using multi-axis force-plates which are not always found in robotic research labs. Here we present a comparison of models using kinematic and kinetic metrics of propulsion to estimate AGRF.

Nine models using measurements of maximum vertical ground reaction force (maxVGRF), vertical ground reaction force at peak AGRF (aVGRF), maximum trailing limb angle (maxTLA), trailing limb angle at peak AGRF (aTLA) and stride length (SL) were used to predict different metrics of propulsion kinetics, including maximum AGRF (maxAGRF), propulsive impulse (PI), maximum AGRF normalized by body-weight (maxAGRFnorm), and normalized PI (PInorm) from participants at speeds [0.6 1.4] m/s. R^2^ and AICc scores were recorded for each model, and the individual participant R^2^ values for the best single and two-factor models for each outcome were examined.

Of the single-factor models, kinematic measurements were the best predictors of the outcome measurements. More specifically, maxAGRF/norm were best predicted by SL (R^2^ = 0.91, 0.82, respectively), and PI/norm were best predicted by maxTLA (R^2^ = 0.84, 0.43, respectively). For the two-factor models, maxAGRFnorm and PInorm were both best predicted by SL and aVGRFnorm, and maxVGRF yeilded the best predictions for maxAGRF and PI. Models predicting maxAGRF/norm better fit individual participants than those predicting PI/norm. These results indicate that maxAGRF can be estimated with reasonable accuracy (R^2^ = 0.92, RMSE of residuals: 1.5% bodyweight, equivalent to a 0.09 m/s increase in velocity) in the absence of a direct measurement of AGRF using both kinematic and kinetic measurements of propulsion.

## I. INTRODUCTION

Propulsion is an important subtask of walking [1]. There are two components of propulsion: push-off posture and propulsive force [2]. Propulsive force is often used as a measure of impairment and recovery post-stroke for individuals with hemiparesis [1]. Hemiparesis is common in post-stroke individuals, and results in a weaker propulsive force from the paretic leg than the non-paretic leg [3]. In other metrics of walking recovery such as gait speed, the recovery of the paretic limb cannot be assessed because patients usually learn to compensate for their weaker leg through less metabolically efficient walking [1], [3]. Thus, metrics that can be separated by leg such as propulsive force are more informative as to the trajectory of a patient’s recovery.

Propulsive force is measured as the positive anterior component of the ground reaction force [4]. The anterior ground reaction force (AGRF) is typically measured on multi-axis force-plates either under treadmill belts or in the ground to observe overground walking [3], [4]. These forceplates are expensive and not always found in robotics labs that focus on exoskeleton research. Additionally, some of these labs may want to test their exoskeletons outside of a traditional lab setting where force-plates are not available. Many have tried to address these concerns by instead turning to the use of inertial measurement units (IMUs) and machine learning algorithms to estimate ground reaction forces [5], [6], [7], [8]. However, IMUs readings may be dependent on the accuracy of placement, unless advanced calibration procedures are used [9], and they can be subject to drift in the signal that must be accounted for. In some applications, it may be easier to measure other components related to propulsion, such as vertical ground reaction force (VGRF) which only needs a one degree of freedom force-plate, trailing limb angle (TLA), or stride length (SL).

These measurements have been shown to be related to propulsive force. Hsaio et al. created a model that described the relationship between TLA, ankle moment, and propulsive force. They were able to show that TLA and ankle moment linearly increased with propulsive force; however, this was performed based on data from only two walking speeds: the participant’s self-selected walking speed and 120% of self-selected walking speed [4]. Similarly, Peterson et al. found that leg extension was positively related to propulsive impulse in post-stroke patients [2]. A study by Lewek and Sawicki evaluated the relationship between multiple definitions of TLA with AGRF in post-stroke patients. They found that for all measures of TLA, there was a positive relationship with peak propulsive force [10]. Martin and Marsh had participants walk at a single speed with 5 different stride lengths. They found that AGRF scaled linearly with stride length [11]. Overall, while there is evidence that these measurements are related to propulsive force, the relationships have not been quantified with respect to a large range of walking speeds, nor with the intent of identifying the ideal model that best predicts propulsion kinetics given a set of available measurements. Testing simple models involving clinically relevant metrics of propulsion may lead to the identification of easily interpretable relationships between these metrics that could enhance our current understanding of walking biomechanics and help identify abnormalities in walking function.

The goal of this research project is to establish whether propulsive force can be estimated accurately using simple linear models in absence of a direct measurement of AGRF, but rather based on the availability of other metrics related to propulsion that are simpler to directly measure, such as VGRF, TLA, or SL, over a wide range of walking speeds.

## II. Methods

### A. Experimental Protocol

A total of 14 healthy young adults participated in this study (5 female, 25 ± 2.66 years old). Participants were asked to walk for one minute at each of five speeds (0.6, 0.8, 1.0, 1.2, and 1.4 m/s), in a randomized order. They were instructed to walk normally at each of these speeds and had a one-minute break between trials. Participants walked on an instrumented split-belt treadmill (Bertec Corp., Colombus OH, USA) and wore four reflective spherical markers; one on each greater trochanter and lateral malleolus (Fig. 1). A ten camera Vicon T40-S passive motion capture system (Oxford Metrics, Oxford, UK) was used to record marker position in 3D space at 500 Hz. Force-plate data was recorded in Simulink (MathWorks Inc., Natick MA, USA) at 500 Hz.

**Fig. 1.**
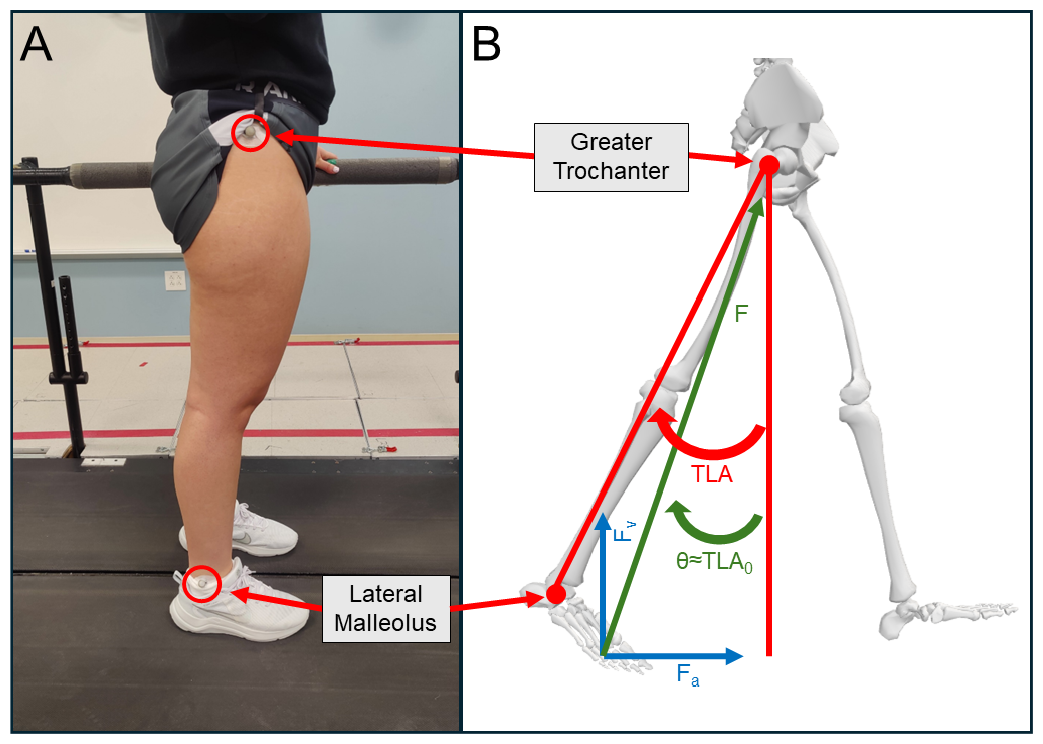
(A) Motion capture marker placement. (B) Free-body diagram of the legs. *F* is the ground reaction force, *F*_*a*_ is the anterior component *F, F*_*v*_ is the vertical component of *F*, and *θ* is the angle between *F* and the vertical. *T LA* is our definition of TLA compared to a previously used definition for TLA (*T LA*_0_) that instead spans from the greater trochanter to the 5th metatarsal head [4].

### B. Mechanics of Propulsive Force

Previous studies have identified critical factors that facilitate forward propulsion: ankle plantarflexion moment [4], leg extension, and trailing limb angle. Peterson defines leg extension as the angle between the laboratory’s vertical axis and the vector from the foot COM to the body’s COM [2]. Trailing limb angle is defined as the angle between the laboratory’s vertical axis and the vector from the greater trochanter to the fifth metatarsal head [12].

Hsiao et al. developed a biomechanics-based mathematical model to quantitatively show the contributions of ankle plantarflexion moment and TLA on peak propulsive force [4]. Instead of using an inverse dynamics approach to calculate ankle moment, the model uses a quasi-static rigid free body diagram approach neglecting dynamic effects on the foot, as well as the weight of the foot. This method has sufficient basis in a study by Wells in 1981 [13], and was validated by Wu and Ladin in 1996 [14]; the results showed that using the projection method of ground reaction force which neglects gravitational and inertial effects on the limbs leads to minimal error in ankle moment using the standard link-segment method.

This section will show that based on this model, a fundamental relationship exists between AGRF, VGRF, and TLA. A free-body diagram of the legs during push-off (Fig. 1B) shows the geometric relationship between *F, F*_*a*_, *F*_*v*_, and *θ*, where *F* is the ground reaction force, *F*_*a*_ is the propulsive force (AGRF), *F*_*v*_ is the vertical component of the ground reaction force, and *θ* is the angle between *F* and the vertical axis.

In general, *F*_*a*_ and *F*_*v*_ are two components of the same vector, so the missing component can be fully reconstructed with knowledge of one component and of the direction of the force vector, i.e.,

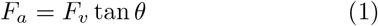

During terminal stance, the vector defining the TLA angle becomes closely aligned with the vector of force extending from the COP at the metatarsals to the COM [10], allowing for TLA_0_ to be substituted for *θ*.

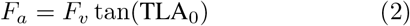

For this study, our lab placed motion capture markers at the ankle instead of the metatarsal to measure TLA. Therefore, a correction factor, *α* is expected, giving a final theoretical relationship of

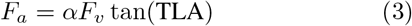

Using a standard least squares model, we estimated the relationship between *F*_*a*_ and *F*_*v*_ *·* tan (TLA) at peak AGRF using data from our experimental protocol. In this model, we also included participant as a random factor. This model had an R^2^ value of 0.81 and AICc score of 58462.1 (Fig. 2), indicating that there is a strong correlation for the relationship described in eqn. (3).

**Fig. 2.**
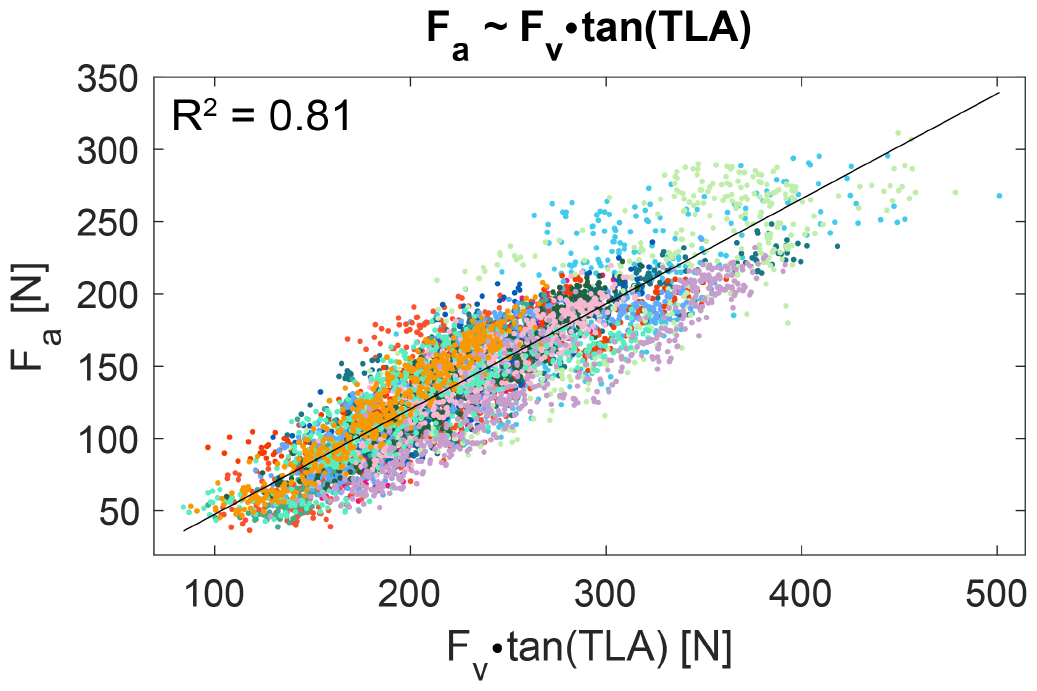
Model representing *F*_*a*_ predicted by *F*_*v*_ *·* tan (TLA). Each participant is represented by a different colored dot. *F*_*a*_ is adjusted by subtracting the best linear unbiased prediction (BLUP) offset for each participant.

In summary, if the direction of the force vector at push-off is known and accurately measured by TLA, we should be able to estimate the anterior component of propulsive force by measuring both *F*_*v*_ and *TLA*. In practice, the accuracy of such an estimation scheme needs to be quantified.

### C. Data Analysis

To synchronize the force-plate and motion capture data, we used QUARC real-time control software (Quanser, Markham ON, Canada) in Simulink to read data from an NI PCIe 6321 card and from Vicon Tracker (Oxford Metrics, Oxford, UK), respectively, in real-time. Force-plate data was low-pass filtered at 150 Hz with a 4th-order zero-shift Butterworth filter. Gaps in marker data were linearly interpolated for a maximum of 25 samples (50 ms). Heel strike and toe off events were defined in post-processing via force-plate data. Heel strike events were defined as the instants at which vertical ground reaction force exceeded 50 N, and toe off events were defined as instants at which vertical ground reaction force dropped below 50 N.

Our outcome measures were peak AGRF (maxAGRF) and propulsive impulse (PI). maxAGRF is defined as the maximum anterior ground reaction force [15]. We used PI as a second outcome measure because PI has been shown to be correlated with center of mass velocity and walking speed [16]. PI is calculated as the area under the positive portion of the anterior ground reaction force [4]. We also evaluated both of these measurements divided by bodyweight (bw) as a second set of outcome measurements (maxAGRFnorm and PInorm) to normalize the data and remove participant-specific biases.

Our predictor measurements in this study were TLA, SL, and VGRF. TLA was defined in two ways for the analysis (Fig. 3). aTLA is the angle between the vertical and the line between the hip and ankle markers at the instant of peak anterior ground reaction force [15]. A second measurement of TLA (maxTLA) was defined to test instances when AGRF is unknown. maxTLA was measured as the maximum angle between the vertical and the line between the hip and ankle markers [17]. Measuring TLA requires the use of a motion capture system which is not available in all labs or when testing outside of a lab setup. Stride length is a kinematic measurement known to be related to TLA [18] that can be measured via center of pressure position on a force-plate or by manually measuring the distance between the start and end location of one’s stride. To calculate SL for this treadmill study, we utilized motion capture data in the following equation:

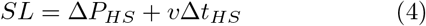

where Δ*P*_*HS*_ is the difference in anterior-posterior position of the ankle marker between the current heel strike (HS) and the previous HS, *v* is the velocity of the treadmill belt, and Δ*t*_*HS*_ is the duration of time between the previous HS and the current HS. For our kinetic predictors, we used vertical ground reaction force defined at two time points (Fig. 3). aVGRF is the value of the vertical ground reaction force at the instant of peak anterior ground reaction force. Again, for instances where AGRF is not available, we measured maxVGRF as the maximum value of the vertical ground reaction force during the time that the anterior ground reaction force is positive (during propulsion). Kinetic measures (aVGRF, maxVGRF) were also normalized by bodyweight (aVGRFnorm, maxVGRFnorm).

**Fig. 3.**
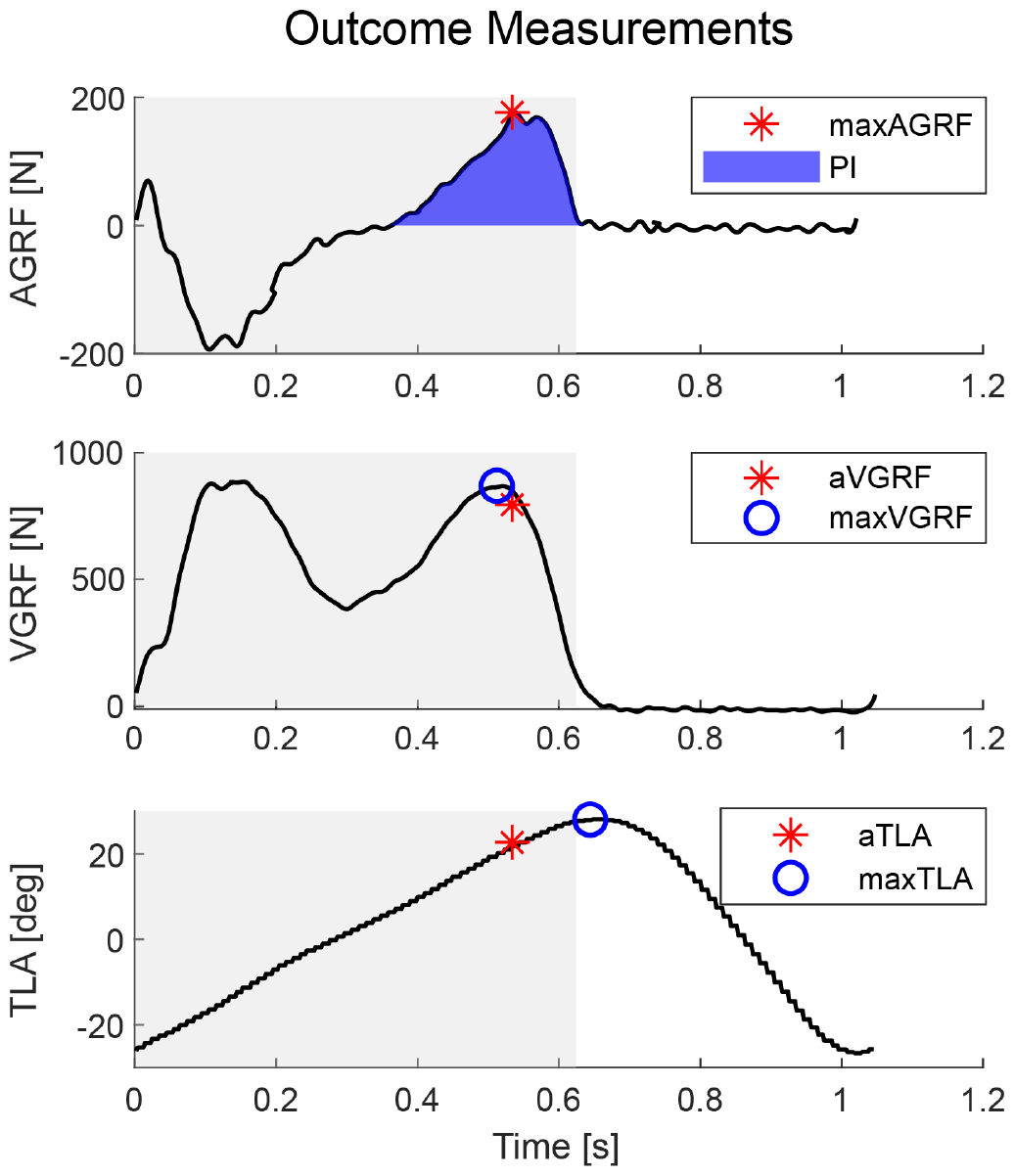
Outcome measurements. Gray shaded region represents duration of stance. (Top) maxAGRF is defined as peak AGRF. PI is the area under the positive portion of AGRF. (Middle) aVGRF is the value of VGRF at the time of maxAGRF. maxVGRF is the maximum value of VGRF during pushoff. (Bottom) aTLA is the value of TLA at the time of maxAGRF. maxTLA is the maximum value of TLA.

**Fig. 4.**
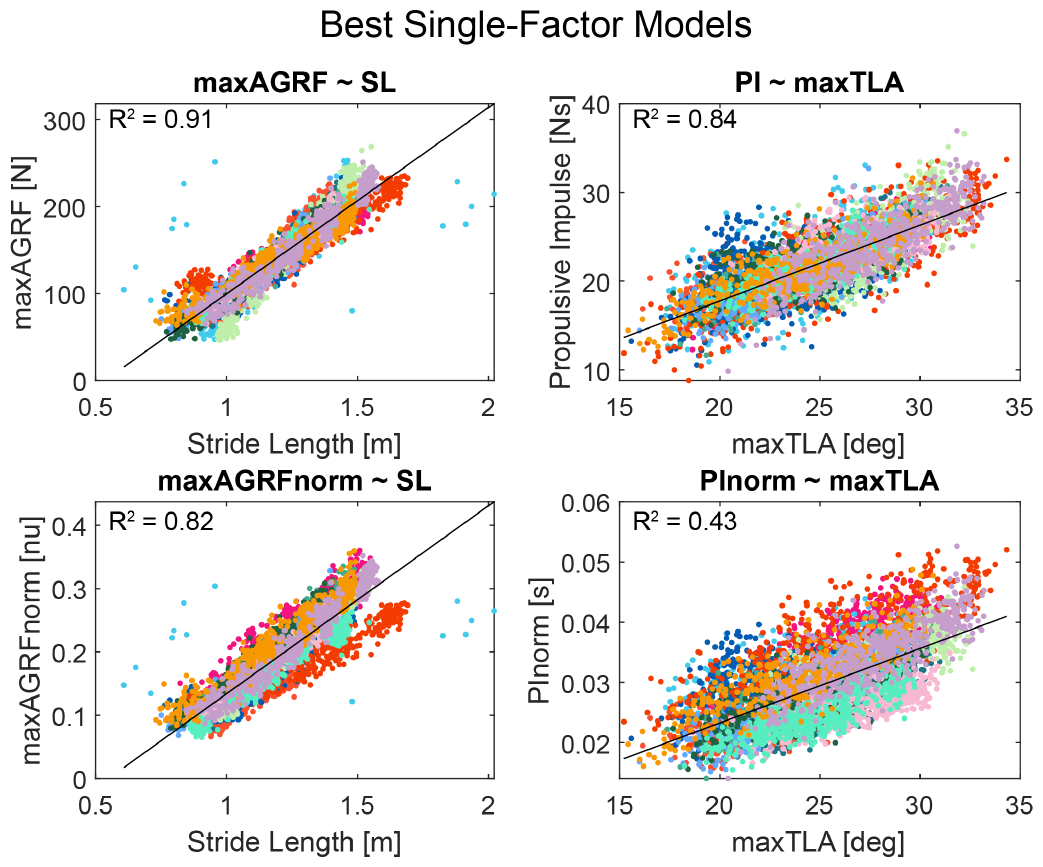
Best single-factor model for each outcome measure. Each participant is represented by a different colored dot. Outcome measure values for the top two figures are adjusted by subtracting the best linear unbiased prediction (BLUP) offset for each participant.

**Fig. 5.**
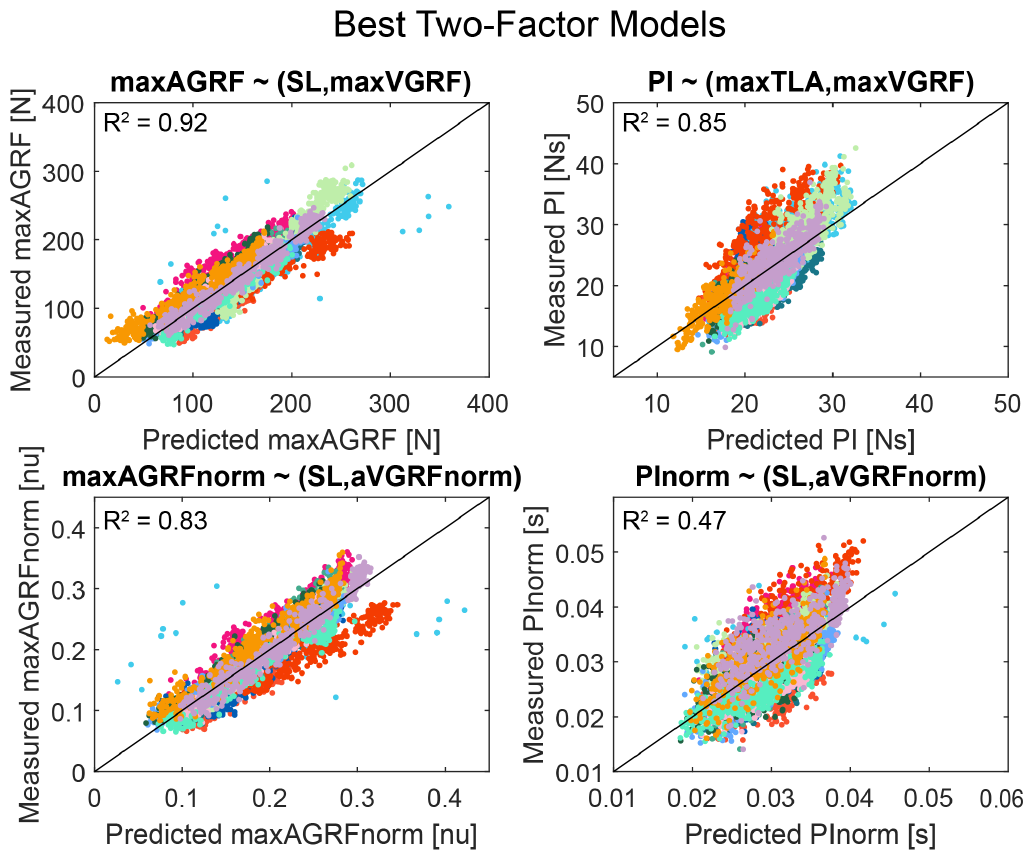
Best two-factor model for each outcome measure. Each participant is represented by a different colored dot. All outcomes were estimated for each stride using the respective prediction equation provided by JMP for that model. Predicted outcome measurements are compared to the actual outcome measurements. The diagonal line represents a 1:1 ratio between predicted and measured outcomes.

**Fig. 6.**
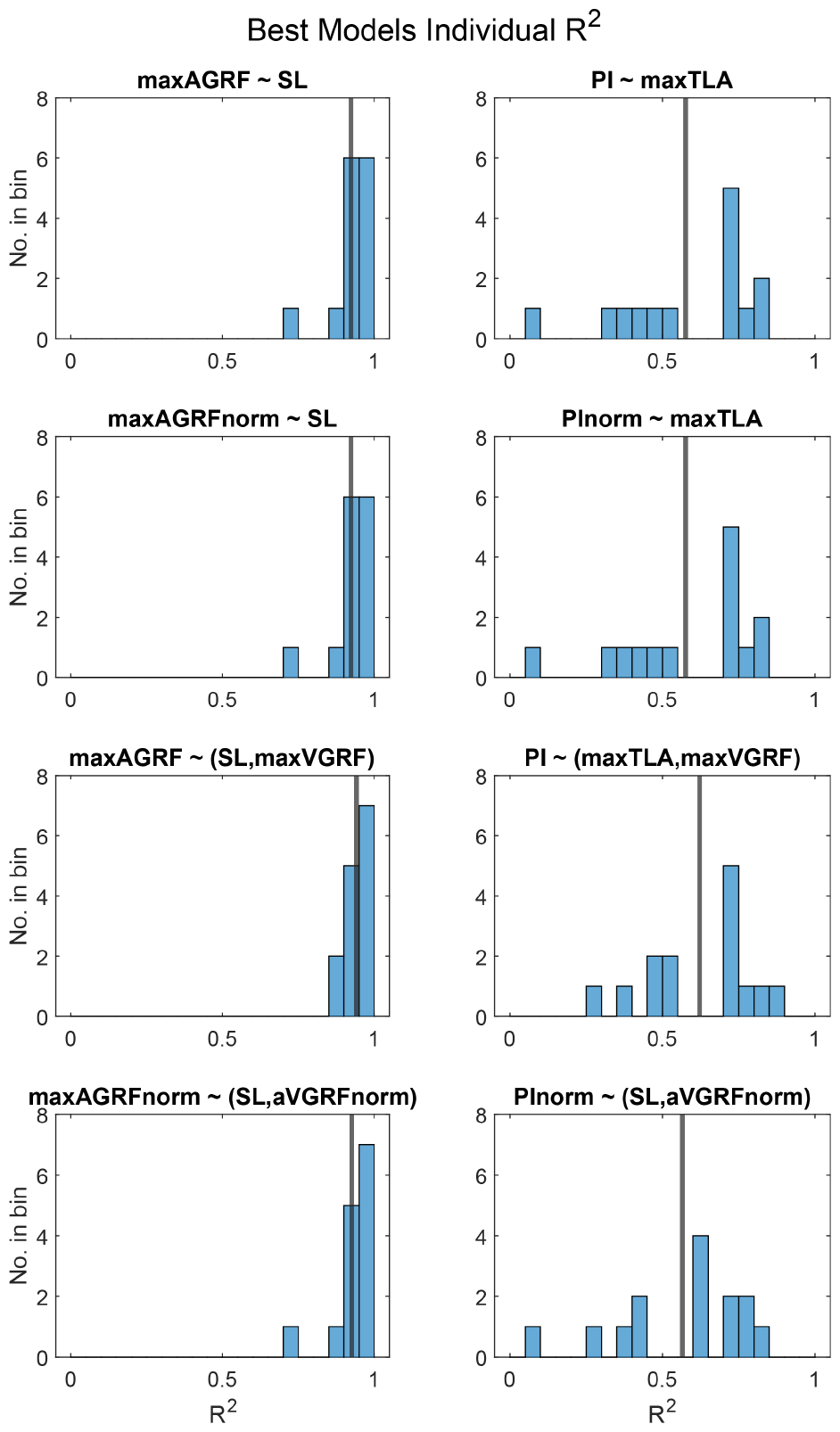
Distribution of individual participant R^2^ values for the best single and two-factor models. The vertical line represents the mean R^2^ value for the distribution.

### D. Statistical Analysis

We used JMP Pro Version 17 (SAS Institute Inc., Cary, NC, USA) to fit a standard least squares model to each of the combinations described in Table 1. We began by finding the relationship between the outcome measures and each of the kinematic (aTLA, maxTLA, SL) and kinetic (aVGRF/norm, maxVGRF/norm) measurements independently to determine which variable explains each outcome the best. Then we tested combinations of kinematic measurements with kinetic measurements (aTLA and aVGRF/norm; maxTLA and maxVGRF/norm; SL and aVGRF/norm; SL and maxVGRF/norm) with the expectation that this would improve the predictions. To estimate a model for maxAGRF and PI, we included participant as a random factor and studied the fixed effects of the predictor(s) of interest. For maxAGRFnorm and PInorm, data from all participants were gathered together under the assumption that between-subject differences were removed when dividing by bodyweight. The kinematic and kinetic factors described in Table 1 were included as fixed effects for each outcome. We recorded R^2^ and AICc values [19] for each model. We then found the R^2^ values for each participant for the best single-factor and two-factor model for each outcome. We found the mean and standard deviation of the individual R^2^ values to understand how well each model could explain individual participants. Finally, the root mean square error (RMSE) of each model was also calculated.

**TABLE I:** Goodness of Fit Results.

| Outcome | Factor 1 | Factor 2 | R<sup>2</sup> | AICc |
| --- | --- | --- | --- | --- |
| maxAGRF | aTLA | - | 0.75 | 60188.8 |
|  | aVGRF | - | 0.37 | 66326.9 |
|  | maxTLA | - | 0.84 | 57584.8 |
|  | maxVGRF | - | 0.69 | 61858.0 |
|  | **SL** | - | **0.91** | **53912.2** |
|  | aTLA | aVGRF | 0.88 | 55668.6 |
|  | maxTLA | maxVGRF | 0.87 | 56016.0 |
|  | SL | aVGRF | 0.91 | 53576.5 |
|  | **SL** | **maxVGRF** | **0.92** | **52761.4** |
| PI | aTLA | - | 0.80 | 30650.6 |
|  | aVGRF | - | 0.71 | 33070.7 |
|  | **maxTLA** | - | **0.84** | **28947.3** |
|  | maxVGRF | - | 0.79 | 30748.7 |
|  | SL | - | 0.83 | 29314.3 |
|  | aTLA | aVGRF | 0.84 | 28937.7 |
|  | **maxTLA** | **maxVGRF** | **0.85** | **28504.4** |
|  | SL | aVGRF | 0.84 | 29103.8 |
|  | **SL** | **maxVGRF** | **0.84** | **28892.9** |
| maxAGRFnorm | aTLA | - | 0.57 | -23159.6 |
|  | aVGRFnorm | - | 0.21 | -19177.8 |
|  | maxTLA | - | 0.71 | -25677.5 |
|  | maxVGRFnorm | - | 0.38 | -20819.3 |
|  | **SL** | - | **0.82** | **-28859.7** |
|  | aTLA | aVGRFnorm | 0.65 | -24563.0 |
|  | maxTLA | maxVGRFnorm | 0.74 | -26491.1 |
|  | **SL** | **aVGRFnorm** | **0.83** | **-29346.3** |
|  | **SL** | **maxVGRFnorm** | **0.83** | **-29161.8** |
| PInorm | aTLA | - | 0.19 | -49101.8 |
|  | aVGRFnorm | - | 0.22 | -49351.3 |
|  | **maxTLA** | - | **0.43** | **-51399.9** |
|  | maxVGRFnorm | - | 0.13 | -48642.2 |
|  | SL | - | 0.42 | -51305.3 |
|  | aTLA | aVGRFnorm | 0.40 | -51127.0 |
|  | maxTLA | maxVGRFnorm | 0.43 | -51398.2 |
|  | **SL** | **aVGRFnorm** | **0.47** | **-51952.1** |
|  | **SL** | **maxVGRFnorm** | **0.42** | **-51323.6** |

## III. RESULTS

Of the single-factor models, kinematic measurements were the best predictors for all outcomes. For models predicting maxAGRF, SL had the highest R^2^ score and lowest (best fit) AICc score (R^2^ = 0.91, AICc = 53912.2). SL was also the best predictor of maxAGRFnorm (R^2^ = 0.82, AICc = - 28859.7). For PI single-factor models, maxTLA had the best R^2^ and AICc scores (R^2^ = 0.84, AICc = 28947.3). maxTLA was also the best predictor for PInorm (R^2^ = 0.43, AICc = -51305.3).

When looking at two-factor models, the best models include a combination of predictors consistent with those identified by single-factor modeling. For maxAGRF, the model including SL and maxVGRF was the best predictor (R^2^ = 0.92, AICc = 52761.4). The best-fit model for maxAGRFnorm was SL and aVGRFnorm (R^2^ = 0.83, AICc = -29346.3). Of all factors predicting PI, maxTLA and maxVGRF had the best fit (R^2^ = 0.85, AICc = 28504.4). Finally, the model including SL and aVGRFnorm was the best fit for PInorm (R^2^ = 0.47, AICc = -51952.1).

At the individual subject level, SL was confirmed to be a strong predictor of maxAGRF (R^2^, mean *±* s.t.d.: 0.92 *±* 0.06). The prediction for the other outcome was less accurate (R^2^ for PI from maxTLA: 0.58 *±* 0.22). Prediction accuracy increased slightly for the two-factor models when estimating maxAGRF (R^2^ when using SL and maxVGRF: 0.94 *±* 0.03, R^2^ when using SL and aVGRF: 0.93 *±* 0.06), and when estimating PI (R^2^ when using maxTLA and maxVGRF: 0.62 *±* 0.17, R^2^ when using SL and aVGRFnorm: 0.57 *±* 0.22).

The model with the highest R^2^ used SL and maxVGRF to predict maxAGRF. For this model, the RMSE was 13.50 N or 1.82 *%* bw. This correlates to an increase of 0.09 m/s in velocity. The next highest R^2^ was the model using SL and aVGRF to predict maxAGRF. For this model, the RMSE was 14.37 N or 1.94 %bw which again correlates to an increase of 0.10 m/s in velocity.

## IV. DISCUSSION

In this study, we evaluated simple linear models that use common gait kinematic and kinetic measurements as predictors of propulsion kinetics, and specifically maxAGRF and PI. This work presents new methods of predicting AGRF useful in conditions where multi-axis force-plates are not available.

We first aimed to determine the best single-factor model for each outcome measure (maxAGRF, maxAGRFnorm, PI, and PInorm). We found that SL was the best predictor for maxAGRF and maxAGRFnorm, and maxTLA was the best predictor for PI and PInorm. This indicates that metrics of propulsion kinematics were the best predictors of propulsion kinetics. For almost all outcomes, maxTLA and maxVGRF resulted in more accurate predictions than aTLA and aVGRF. Based on eqn. (3), we expected that aTLA and aVGRF would be better predictors than their counterparts because they occur at the same time as the outcome measure (maxAGRF). This means that knowledge of the timing of AGRF is not necessary to predict this quantity, which indirectly supports the possibility of estimating it in absence of direct measurements.

For the two-factor models, we saw similar results to the single-factor models, except with maxVGRF as a second factor. The inclusion of a second factor improved the R^2^ value which was expected, but these models also had lower AICc scores for their outcome measure, which suggests that these additional measurements indeed provided new information content in the model, and were not a consequence of overfitting the model with too many parameters. For maxAGRFnorm and PInorm, the models using SL and aVGRFnorm had a higher R^2^ and lower AICc score than the models using SL and maxVGRFnorm; however, there is a small difference between the two, implying that the kinematic factor (SL) governs the fit of the model whereas the kinetic factor is not as important. Additionally, with the exception of the models predicting PInorm, the single and two-factor models all had better R^2^ and AICc values than our theoretical model that used *F*_*v*_ *·* tan (TLA) (or aVGRF *·* tan (aTLA)) to predict *F*_*a*_ (or maxAGRF).

The individual R^2^ values had the same distribution between the models for maxAGRF and maxAGRFnorm, and the models for PI and PInorm. Because the difference between each outcome and its normalized equivalent is a factor of bodyweight, and each outcome/normalized outcome pair is best predicted by the same prediction measurement, it is expected that these distributions would be identical, despite their R^2^ values being different. maxAGRF/norm better predicted individual subjects than PI/norm which had a minimum participant R^2^ of 0.06. Of the two-factor models, the model predicting maxAGRF best fit individual subjects, followed closely by the model predicting maxAGRFnorm. All models predicting maxAGRF/norm performed similarly as did those predicting PI/norm.

Hsiao et al. found that increases in TLA accounted for 65% of the increase in peak AGRF as opposed to 33.7% contribution from an increase in ankle moment [4]. In our study we found that maxTLA best predicted PI; however, SL best predicted maxAGRF. It is possible that the TLA data we measured had more noise than SL did. We removed strides with poor motion capture quality such as artifacts or missing marker data which resulted in removing 8.01 *±* 9.25% of strides per participant. Though these strides were removed for all outcome measures to keep the sample size consistent, the fact that these many strides were removed due to poor data quality surrounding maxTLA could explain why TLA was not the best predictor for maxAGRF even though it was in the Hsiao et al. paper.

IMUs are another method of estimating AGRF that is being investigated. Miyazaki et al. fixed IMUs to the thorax and lumbar spine of healthy adults to find the correlation between PInorm and change in velocity of the IMU [20]. Their best correlation was from the change in velocity of the lumbar IMU in their “lean-slow” condition (*r* = 0.89, *p <* 0.001). In the “lean-slow” condition, subjects leaned forward during right heel strike while they walked at a speed less than their preferred speed, to mimic a pattern of walking commonly found in stroke patients. Most of their correlations for PInorm were high, mostly ranging from *r* = 0.75 to *r* = 0.89. Though the numbers cannot be directly compared, we also had high correlations of PI (R^2^ = 0.71 to R^2^ = 0.85), indicating that we are able to predict PI similarly.

Revi et al. used IMUs to predict peak propulsion force and propulsion impulse in both healthy and post-stroke individuals [8]. Here, IMUs were placed on the shank, thigh, and pelvis. In their healthy cohort, their model was able to achieve an RMSE of 2.37 %bw in their validation set for maxAGRFnorm. The RMSE for our best maxAGRF model is 1.82 %bw. Their model for estimating PI in their validation set achieved an RMSE that we calculated to be equivalent to 0.31 %bw*·*s (assuming a duration of positive propulsion of about 360 ms). The RMSE for our best PI model is 0.29 %bw*·*s. Based on the RMSE scores, both of our models performed better than those presented in Revi et al. One limitation of our study is that we only tested on healthy young adults. AGRF is an important metric in post-stroke rehabilitation research. In order to apply the techniques described in this paper, an additional study should first be performed using post-stroke individuals to derive the equations of fit that population.

We were able to show that AGRF can be predicted by kinematic and kinetic measurements of gait. More specifically, of single-factor predictors, kinematic factors best predicted outcomes related to propulsive force; maxAGRF/norm is best predicted by SL and PI/norm is best predicted by maxTLA. For two-factor models, maxAGRF and PI both added the use of maxVGRF to their single-factor model, and maxAGRFnorm and PInorm added aVGRFnorm. There was no strong correlation for the models predicting PInorm. Future work will apply these models to post-stroke individuals.

